# Recovering ecological interactions by mining non-target data from whole genome re-sequencing projects

**DOI:** 10.1101/2025.01.17.633498

**Authors:** Mirkka Jones, Pasi Rastas, J. Camilo Chacón-Duque, Michelle F. DiLeo, Abhilash Nair, Vicencio Oostra, Marjo Saastamoinen, Anne Duplouy

## Abstract

The study of parasitic species can shed light on aspects of their host’s ecology. Such interactions are however often unknown or understudied due to the difficulty to detect and/or quantify many infections. Whole-genome sequencing and re-sequencing data have been generated at an increasing rate and reduced costs over the last two decades. Projects based on whole organisms, like whole insect specimens, provide genomic material for the target taxon, but may also include sequencing reads from associated microbes and other parasites. Here, we screened for the presence of non-host reads in a collection of whole-genome (re-) sequence projects from the Glanville fritillary butterfly, *Melitaea cinxia,* a model organism in research on the ecology and evolution of species in spatially structured and fragmented landscapes. We identified infections with different bacteria and eukaryotic parasites, which are shared between populations and underly both previously known and new biotic interactions for this butterfly species. The bacterial symbiont *Wolbachia*, usually common in insects, was found at relatively low prevalence, while *Spiroplasma* was ubiquitous across samples from several European populations of the butterfly. Additionally, we confirmed expected rates of larval parasitism by at least two parasitoid wasps. Such results provide proof of principle that key ecological interactions can be uncovered efficiently from existing data, an important first step to characterising the role host-associated organisms play in shaping the ecology and evolutionary history of their host species.

## Introduction

Whether costly or beneficial, infections can have large impacts on their hosts’ life-histories, population dynamics and genetic diversity (Glasl et al. 2016; Hochachka et al. 2021; Rogowski et al. 2020; Tollenaere et al. 2014). For example, In the Åland Islands, the infection of 10% or more of all available *Melitaea cinxia* butterfly larvae by the parasitoid wasp *Cotesia melitaearum* can severely affect local butterfly host populations (Lei et al. 1997a; Shaw et al. 2009), while neighbouring populations might remain unaffected by the parasitoid (Lei et al. 1997b; van Nouhuys 2016). Similarly, bacterial symbionts that are maternally transmitted, such as *Wolbachia* and *Spiroplasma*, can manipulate diverse host life-histories and other reproductive traits (Correa & Ballard 2016; Duplouy & Hornett 2018). They can, for example, induce cytoplasmic incompatibility (CI) between infected males and uninfected females (Charlat et al. 2006; Hornett et al. 2008; Koehncke et al. 2009; Pollmann et al. 2022), or protect their host against pathogens (Teixeira et al. 2008; Łukasik et al. 2013), thus boosting the reproductive output of the symbiont-infected females compared to uninfected ones. These manipulations facilitate the spread of the symbiont through the host population, and in turn lead to a reduction in the genetic diversity of their host, as host infected mitochondrial haplotypes hitchhike successful infections (Charlat et al. 2009; Deng et al. 2021; Graham & Wilson 2012). Such infections are, however, not always included in ecological and evolutionary studies of host species, as they often remain hidden because they may be tricky to detect and not always abundant across their host geographical distribution.

The development of next-generation sequencing methods has allowed the birth of metagenomics, which aims to sequence and analyse the reads from both a targeted host and its diverse associated organisms. In insects, metagenomic analyses from whole-body sequencing projects have, for example, identified a wide diversity of previously hidden, or difficult to study or culture bacterial endosymbionts (*i.e*., microbes that live inside host cells or tissues)(Gibson et al. 2014; Twort et al. 2022; Vancaester & Blaxter 2023). These studies have revealed key ecological and evolutionary species-interactions (Chellappan & Ranjith 2021; Liang et al. 2020; Vancaester & Blaxter 2023), and provide insights into many complex species-interactions, including the taxonomic identity and genetic diversity of associated species, their prevalence and biogeography, as well as their function (Ghanavi et al. 2021; Susi et al. 2019; Zhang et al. 2018). Nonetheless, despite the dramatic effects that infections can have on their hosts, and despite the development of metagenomic approaches, whole-genome projects rarely include analyses investigating the metagenome of the targeted host species. Consequently, many infections remain uncharacterized, significantly limiting the conclusions drawn from studies of their host populations.

The Glanville fritillary butterfly, *Melitaea cinxia*, is a well-known model-species for the study of the eco-evolutionary consequences of habitat fragmentation on wild species (Duplouy et al. 2013; Hanski & Heino 2003; Kahilainen et al. 2018). The species is widespread across Eurasia, where it is known to feed on a few host plant species (Hanski et al. 2004), including the ribwort plantain, *Plantago lanceolata*. In particular, a metapopulation of the butterfly in the Åland Islands, an archipelago between the coast of Finland and Sweden in the Baltic Sea, has been the focus of 30 years of ecological and evolutionary dynamics studies (Ojanen et al. 2013; Saastamoinen 2022). Studies of this butterfly metapopulation have been complemented by studies of their associated community of specialized parasitoid wasps (Lei et al. 1997a; van Nouhuys et al. 2012), as well as a handful of recent microbiota studies (Duplouy et al. 2018; Duplouy et al. 2020a; Minard et al. 2019; Rosa et al. 2019). The rearing of field-collected *M. cinxia* larvae in the laboratory shows that parasitoid wasps emerge each year from about 30% of the larvae (Montovan et al. 2015), and that the presence of a parasitoid infection affects the species composition of the larval gut microbiota (Minard et al. 2019). Although these results have provided insights on trophic and symbiotic interactions within this ecological system (Lei et al. 1997a; van Nouhuys et al. 2012), several aspects of these interactions remain unclear. For example, it is unclear whether all parasitism attempts are successful, or whether the parasitoid emergence rate only represents a fraction of the parasitoid eggs laid.

The exceptionally detailed understanding of the ecology of *M. cinxia* has recently been complemented with a large amount of genetic and genomic information, including three high-quality versions of the whole genome and chromosomal assembly of the butterfly species (Ahola et al. 2014; Smolander et al. 2022; Vila et al. 2021), and ongoing population re-sequencing studies. Here, we analyse several *M. cinxia* genomic datasets as a source of non-target genomic material reflecting diverse prokaryotic and eukaryotic infections in larvae and adult butterflies from the Åland Islands (Finland), as well as from adults collected across Europe and North-Africa (France, Morocco, Spain, Switzerland). Specifically, we investigate presence, prevalence and geographical range of an array of parasites, including the common endosymbiotic bacteria *Wolbachia* and *Spiroplasma*, and the *Cotesia* and *Hyposoter* larval parasitoids. The detection of microbial symbiont species provides the foundation for further characterizing how they might affect the population dynamics and genetics of their host species. Detected parasitoid infection rates, in turn, provide information on both the prevalence and the potential amplitude of their impacts on butterfly host populations.

## Materials and Methods

### a) Origin of raw sequence data

The Glanville fritillary butterfly, *Melitaea cinxia* (Linnaeus, 1758), is a widespread checkerspot species across Europe, and the model organism for 30 years of ecological and evolutionary studies on the effects of habitat fragmentation in the Åland Islands, an archipelago in the Baltic Sea region (Ojanen et al. 2013). Across the butterfly’s geographical range, the species inhabits meadow habitats, where larvae feed on few host plants, including *Plantago lanceolata* (Linnaeus, 1753) and *Veronica spicata* (Linnaeus, 1753). The butterfly’s eggs, larvae and pupae are parasitised by several common parasitoid wasp species, including specialized species such as *Hyposoter horticola* (Gravenhorst, 1829) and *Cotesia melitaearum* (Wilkinson, 1937). These wasps belong to the Braconidae and Ichneumonidae groups, respectively, and are known to have important impacts on the butterfly’s population dynamics (Lei et al. 1997a). The bacterial microbiota associated with larvae collected in the field in the Åland islands, or reared in the laboratory, have been previously characterized through *16S* rRNA gene metabarcoding approaches (Duplouy et al. 2020a; Minard et al. 2019).

The first assembly of the genome of *M. cinxia* was released in 2014 (Ahola et al. 2014), followed by two chromosome-level assemblies in 2021 (Vila et al. 2021) and 2022 (Smolander et al. 2022). We retrieved the raw sequence data from five whole genome sequence datasets, including two genomic assembly projects (Ahola et al. 2014; Smolander et al. 2022) with NCBI references #[PRJNA191594](https://www.ncbi.nlm.nih.gov/bioproject/191594) and #PRJNA607899, respectively, and from three additional independent projects comparing genomic characteristics of different local or regional populations (labelled Adults Europe #C; Åland larvae & adults #D; and Åland larvae #E). Dataset #D was derived from both larval and adult specimens collected in the Åland Islands in 2009 and 2010, dataset #E from wild larval samples collected in the Åland Islands in 2018 and 2019, while dataset #C included only wild adult specimens collected during summer 2020 in France, Morocco, Spain, and Switzerland. The sample preparation and sequencing techniques vary between projects, including NovaSeq, NovaSeq 150bp Paired-Ends, and NextSeq 170bp PE (as described in Table 1).

**Table 1:**
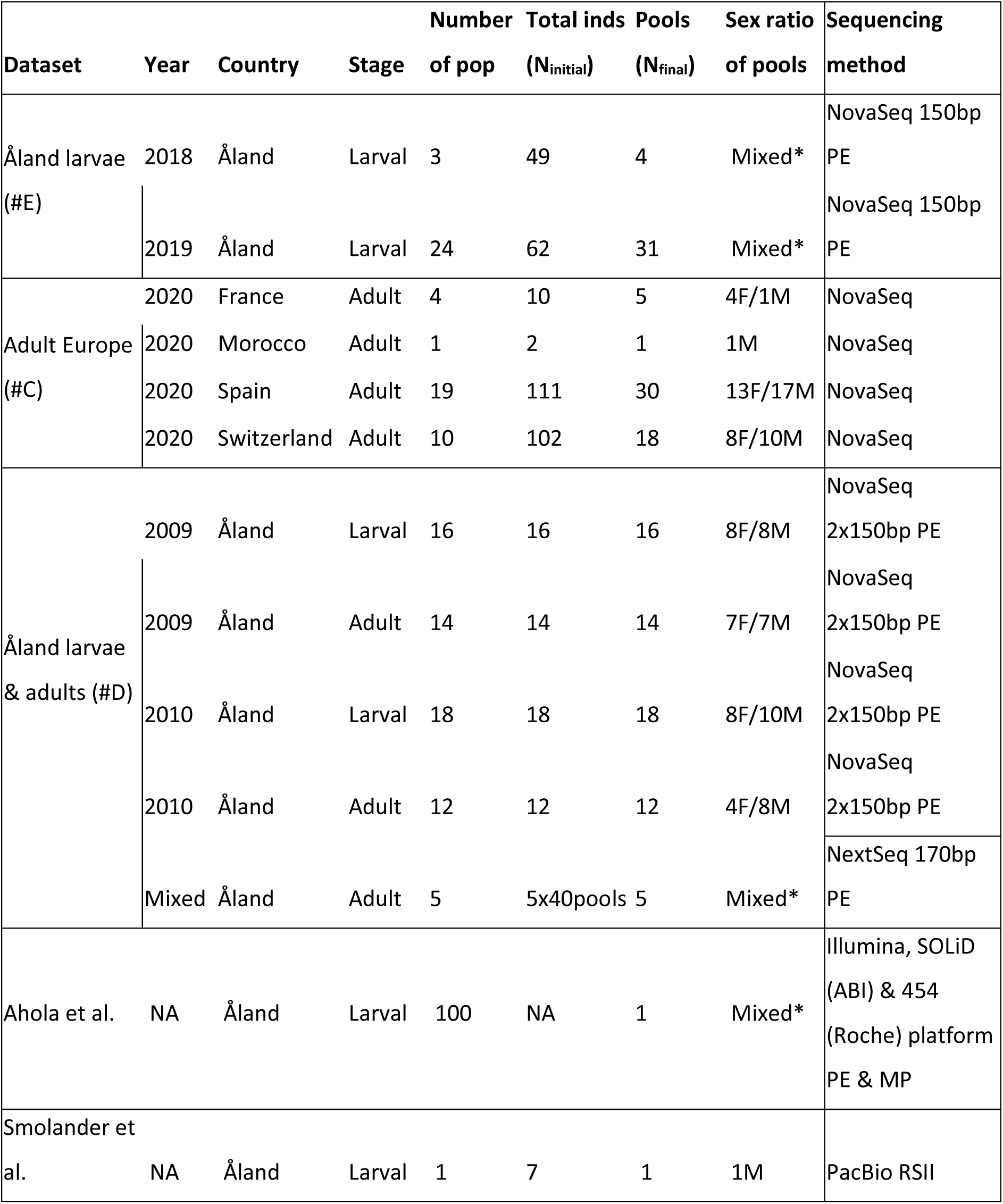
Sample size, country of origin, developmental stage, and sex for each of the genomic project screened for non-host reads. The * samples were pooled without prior knowledge of the sex of the larvae.

Unfortunately, the low coverage for each specimen in all projects compromised the deep characterization of the metagenome of each specimen individually. Thus, to improve our ability to detect non-host material, we pooled all samples from the same dataset, same population and same sex (when known) together. This reduced the total number of samples from the original N_initial_=425 specimens (plus the two full genome projects) to N_final_=166 samples (plus the two full genome projects). Details on the pooled sample size, country of origin, life stage and sex of the samples, as well as the sequencing method used for each project, are described in Table 1.

### b) Sex characterization of larval samples

Although we characterized the sex of the adult specimens through visual inspection of their external genitalia prior to dissections, the larvae of this butterfly species do not show any external sexually dimorphic traits, and thus their sex could only be determined after sequencing. Females were identified based on homozygosity across a set of 22 Z-chromosome specific single-nucleotide polymorphism loci (Kahilainen et al. 2022).

### c) Screening for non-host genomic material

We trimmed the Illumina paired-end reads to remove Illumina adapter sequences and any low-quality bases from the start and end of reads using Trimmomatic version 0.39 (Bolger et al. 2014) with the ILLUMINACLIP adapter-clipping settings 2:30:10 LEADING:3 TRAILING:3 SLIDINGWINDOW:4:15 MINLEN:50. We then used the SPAdes assembly toolkit version 3.15.0 (Bankevich et al. 2012) (https://github.com/ablab/spades) with the --meta (metagenomic) flag to denovo assemble the trimmed paired reads into contigs (Nurk et al. 2017). To reduce heterozygosity, we applied the Redundans pipeline (Pryszcz & Gabaldón 2016) to the assembled scaffolds, and later annotated them to NCBI TaxIDs using BLASTN searches (BLAST v. 2.12.0, https://docs.csc.fi/apps/blast/) against the NCBI non-redundant nucleotide database, retaining one aligned contig per scaffold (max_target_seqs 1). We only saved the best alignment for each query-subject pair (max_hsps 1), and with an E-value less than 1 × e−25. We then used BWA-MEM (Li & Durbin 2009; Li & Durbin 2010) to map all the original trimmed and corrected reads to the reference scaffolds and converted the results into sorted bam format files containing sample, sequence and mapped read data with SAMtools (Li & Durbin 2009). Finally, we generated BlobPlots and associated statistics at the genus taxonomic rank for each assembly with Blobtools v1.1 (Laetsch & Blaxter 2017) based on the BLASTn similarity search results, and using coverage and GC content values to differentiate host genomic material (Table S1).

Contamination with any non-host material was considered real, when the value of the ‘bam0_read_map_p’ output from the Blobtools analysis was above 0.5%. Three samples (from #D), were removed prior to subsequent analyses because of their overall lack of non-host data (all non-host hits <0.5% in AF10-173-6, AF10-7007-1, and 09-8544-001-1-5). Additionally, to check that our results were repeatable, we compared the outputs of our Blobtools analyses with the outputs from CC-Metagen analyses (https://github.com/vrmarcelino/CCMetagen)(Clausen et al. 2018; Marcelino et al. 2020) for five randomly chosen samples (AF18_2155_1, AF19_48_1, AF19_57_1, AF10-13341-3 and AF19_1956_1).

### d) Non-host community composition analyses

By combining the Blobtools results at the genus level from all the samples, and only selecting the non-host species with a total ‘bam0_read_map_p’ output equal to or above 1%, we created a community matrix of 78 non-host species. This included prokaryote species, as well as some eukaryote species. Hits to Diptera taxa and to *Homo sapiens* were considered to represent clear sample preparation contaminations and were manually removed before any further analyses.

We tested for dissimilarities between treatments (Country, Dataset or Life stage) in this community matrix, using the ANOSIM statistic R test with N=9999 permutations in the VEGAN package in R v.2023.12.0+369 (RCoreTeam 2020).

### e) Building genome assemblies of common parasites and symbionts

We retrieved a representative list of high-quality *Wolbachia* and *Spiroplasma* genomes from the NCBI Genome Data Hub (by April 2022), to build independent reference databases of each microbial symbiont (Valerio et al. 2024a). The available NCBI genome projects were filtered to only include ‘Complete’ and/or ‘Reference’ genomes. Similarly, we built a reference database for the parasitoid wasp *Cotesia sp.* using only the two high-quality genomic projects available from NCBI (by April 2022), including the genome assembly of *C. congregata* (WGS project #CAJNRD03), and the genome assembly of *C. glomerata* (WGS project #JAHXZJ01).

In an attempt to build the *Wolbachia, Spiroplasma* and *Cotesia* assemblies from our metagenomic analyses, we combined the reference *Wolbachia*, *Spiroplasma* and *Cotesia* genomes and two *M. cinxia* full genomes (Smolander et al. 2022; Vila et al. 2021) into a single fasta file, after renaming their sequences with a prefix Wo, Sp, Co, or ci, respectively. We used BWA-MEM (Li & Durbin 2009; Li & Durbin 2010) to map raw reads from all other projects to this multi species reference. Reads with an alignment score >= 240 (max score is 300, 150bp+150bp reads) were then assembled using the SPAdes software (Bankevich et al. 2012), following the same protocol as in the Blobtools pipeline described above.

## Results

### a) Detection of infections from genomic material of *M. cinxia*

Despite combining samples from the same population, year and sex (when possible), the depth of the non-*M. cinxia* genomic data and total amount of reads assigned to *M. cinxia* and non-*cinxia* organism per sample remained low for many samples (Table S2). The amount of non-host material from both host genome sequencing projects (Ahola et al. 2014; Smolander et al. 2022) were too low to be confidently further examined (data not shown). The number of reads assigned to *M. cinxia* from the three re-sequencing projects varies from 7485 to 44687900 reads per sample (with an average=20540118.3), while the number of reads assigned to non-host varies from 4024 to 43524459 reads per sample (with an average=707841) (Table S2). Screening of these samples still provided information on several expected and unexpected infection patterns. The outputs of the Blobtools analyses were similar to those of CC-Metagen analyses on five replicated samples (Table S3). The larval samples AF18_2155_1 and AF19_1956_1 were poor in non-host material, while sample AF19_48_1 contained *Spiroplasma* reads, sample AF19_57_1 contained genomic material of Braconidae wasps and their polydnavirus, and sample AF10-13341-3 included reads from *Pseudomonas*, *Enterobacter*, *Klebsiella*, and *Gluconobacter*.

The analysis of dissimilarity between the non-host communities associated with our samples indicated that both the source dataset (ANOSIM, R=0.124, p=1e-4, Figure 1A), and butterfly life stage (ANOSIM, R=0.0222, p=0.0077, Figure 1B) affected community composition. Country, however, had no significant effect on the composition of non-host communities (ANOSIM, R=-0.128, p=0.996, Figure 1C). Our analyses, however, only investigate the non-host species community at the genus level, and analyses at the strain level could show different community patterns between countries.

**Figure 1:**
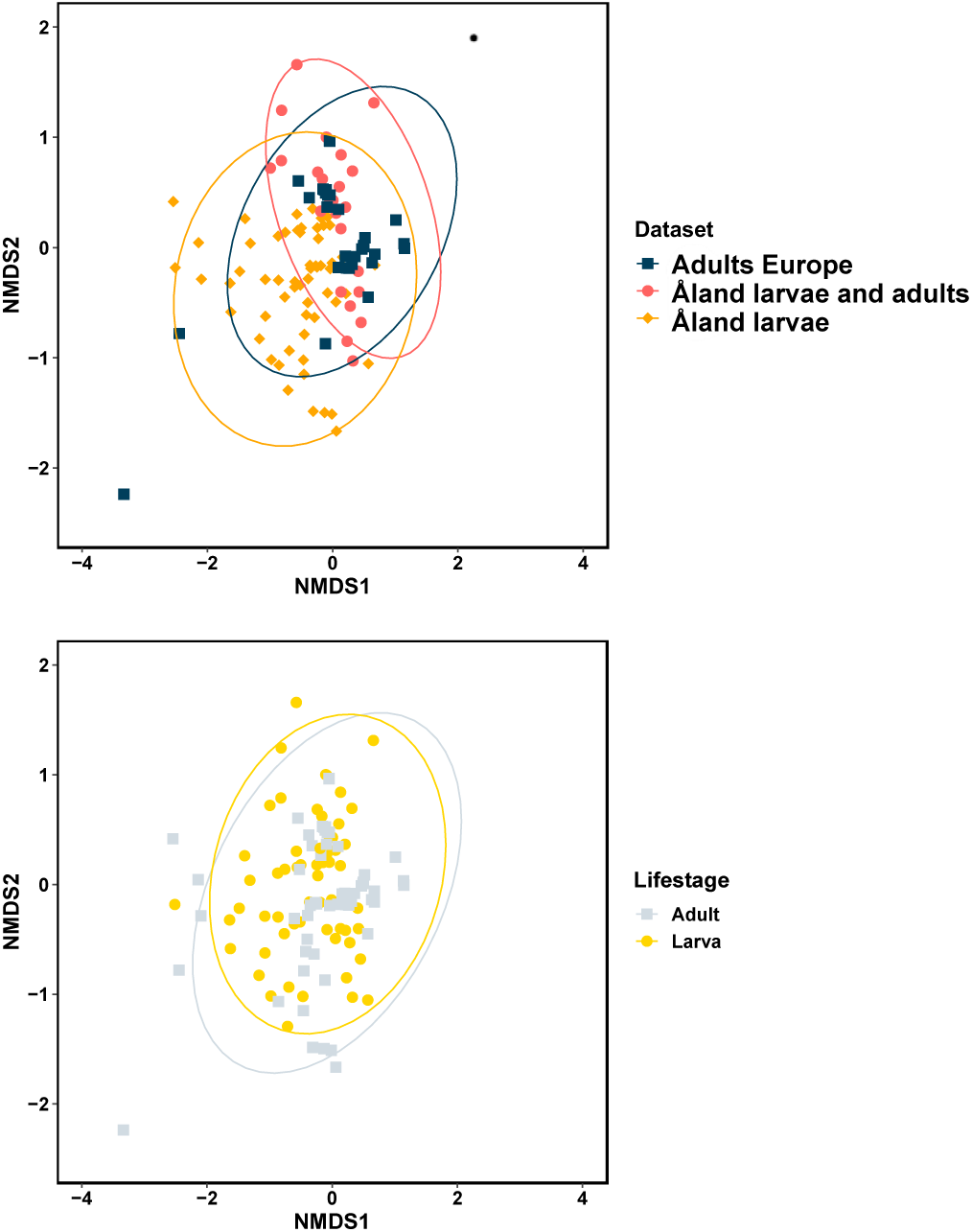
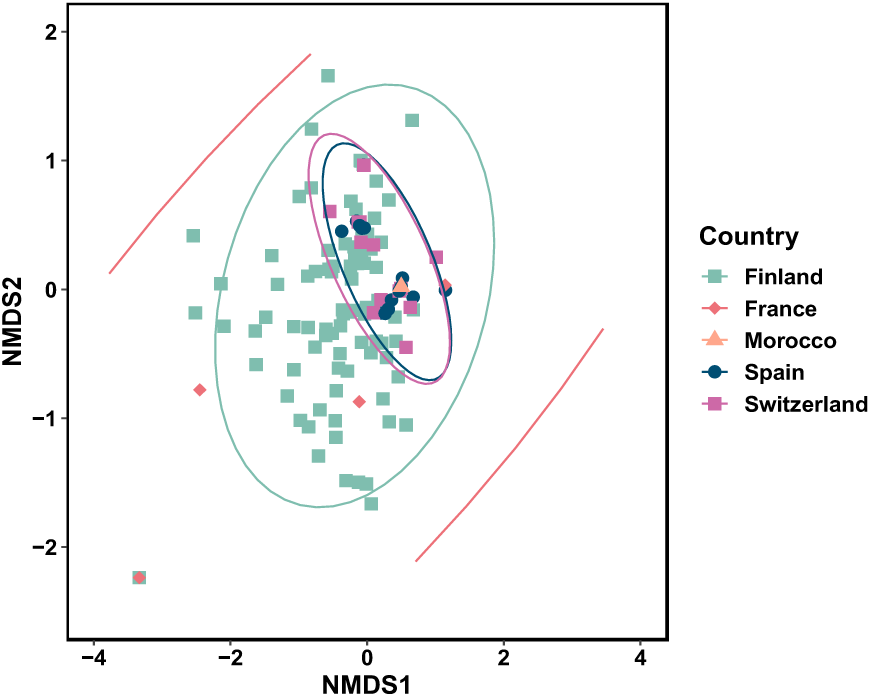
NMDS1 graph of the non-host community composition. Samples are coloured as (Top) per dataset, (Middle) per life stage, and (Bottom) per country of origin (Salmon ellipse for France covers all other ellipse, and is only shown partially in the graph; The Moroccan population only includes 1 sample and is not represented by any ellipse).

Our analyses showed that four and six *M. cinxia* samples were contaminated with genomic material from two genera of parasitoid wasps: *Cotesia sp.* and *Hyposoter sp.*, respectively (Duplouy et al. 2015; van Nouhuys 2016). Such contamination was found in both female and male larvae, but never from adult specimens. Genomic reads (ie. N=186432 reads or above) from the braconid *Cotesia sp*. wasp, as well as from the wasp-associated bracovirus (Benoist et al. 2017; Savary et al. 1999), were identified in four larval samples, from the Eckerö and Hammarland communes in Åland (patch #572, 1646, 57, and 1056). Genomic reads from the Ichneumonid *Hyposoter sp.* wasp, as well as reads from the wasp-associated ichnovirus (Clavijo et al. 2011; Dorémus et al. 2013), were detected from six larvae (Table S1) collected from the communes of Saltvik, Hammarland and Sund in the Åland Islands (patch #123, 535×2, 9620, 9025, 9051).

We also detected genomic material from the bacterial endosymbionts *Wolbachia*, *Spiroplasma*, and *Rickettsiella*, which are generally ubiquitous in insects (Duplouy & Hornett 2018; Raina et al. 2018; Weinert et al. 2015). *Spiroplasma* infections were encountered in 23/86 (26.7%) larval and 22/83 (26.5%) of adult host samples, from both female and male specimens. *Spiroplasma* was detected from specimens from all source countries, except from Morocco, which is only represented by one sample in our dataset. Despite its high prevalence in our samples, *Spiroplasma* always represented less than 10% of the total reads per sample (ie. 9400 or more reads/sample). In contrast, *Wolbachia* was represented by 3.7% to 43.4% of the non-host reads per sample (155526 or more reads/sample), and occurred at a lower prevalence in our datasets, with only five adult samples found infected (5/166, 3%) (Table S1). These *Wolbachia*-infected samples include three pools of females and two of males, from either Spain or Switzerland. The other endosymbiotic bacterium *Rickettsiella* was detected in a unique larval sample, and was represented by 54% of the non-host reads of that sample. Many other bacterial species were also found sporadically, including *Erwinia* (one French adult and one Finnish larval samples, 7%, and 80% of the non-host reads, respectively), *Rhanella* (two larval samples, 5% and 58% of the non-host reads), *Serratia* (one French adult sample, 63.5% of the non-host reads), and *Yersinia* (one adult and five larval samples from the Åland Islands, 1.6% to 50.4% of the non-host reads respectively). One adult sample from the commune of Vall d’Estos in Spain was highly dissimilar to all others (Table S1). This Spanish sample was almost fully contaminated with the bacterium *Proteus* (96.9% of the non-host reads), and it was the only sample carrying this bacterium in our dataset. This sample was not excluded from the analyses but was excluded as a visual outlier from Figure 1. Four microbial genera, *Gluconobacter, Enterobacter, Actobacter* and *Alphabaculovirus* were only found in the adult and larval Åland dataset #D, while all other microbes were identified from several datasets.

### b) Genomic assemblies of identified non-host organisms

By combining all reads from Åland, we were only able to assemble a 11.2Mb genomic assembly of *Cotesia* (eg. expected whole genome size >180Mb), and a 65Kb genomic assembly of *Spiroplasma* (ie. expected whole genome size ≈1.2Mb). We also assembled a 95Kb assembly of *Wolbachia* genome, by combining reads from the five infected specimens (ie. expected whole genome size ≈1.2Mb, Table 2). The covered regions spanned several host genomic scaffolds, and each at a very low abundance (Figure S1). We were unable to retrieve and compare sequences of commonly characterized loci of *Cotesia*, *Spiroplasma* or *Wolbachia* strains to our partial assemblies (Duplouy et al. 2015; Kankare et al. 2005). The comparison of the largest contigs from each partial assembly against the NCBI nucleotide database (Blastn search conducted in December 2024, Table 2) however supports the assigned identities of our three partial assemblies. Notably, although the GC contents of both the *Cotesia* and *Spiroplasma* partial assemblies correlate with previous genomic analyses on these organisms (36% and 25%, respectively), our *Wolbachia* partial assembly exhibits a much higher GC content than other *Wolbachia* genomic projects (49% instead of the usual 34%) (Table 2).

**Table 2:** Statistics and features of three partial genomic assemblies, and best match output (i.e. blastn) between the largest contig of each assembly and the nucleotide NCBI database (December 2024).

| Organism | Number of contigs | Size | Smallest contig | Largest contig | Mean contig | GC % | Largest contig's best Blastn hit (NCBI, Dec 2024) |
| --- | --- | --- | --- | --- | --- | --- | --- |
| *Cotesia sp.* | N=73194 | 11212 KB | 122 bp | 6096 bp | 471.4 bp | 38.3 | *Cotesia congregata*<br>(#FM212914) |
| *Spiroplasma* | N=59 | 65 KB | 184 bp | 5050 bp | 915.8 bp | 26.2 | *Spiroplasma* of *Seladonia*<br>*tumulorum* (#OZ026545) |
| *Wolbachia* | N=112 | 95 KB | 130 bp | 2364 bp | 653.5 bp | 48.9 | *Wolbachia* of *Drosophila*<br>*pseudotakahashii* (#CP110359) |

## Discussion

Our screening for non-host genomic material in five independent whole genome sequencing datasets of the butterfly *M. cinxia* revealed the presence of genomic material from different non-host micro- and macro-organisms. Our approach of exploring the non-host genomic material from whole genome sequencing projects for evidence of other species is not novel, and has previously provided insights into a hidden diversity of microbes associated with diverse insect species (Ghanavi et al. 2021; Twort et al. 2022; Vancaester & Blaxter 2023). However, this is, to our knowledge, the first time that the method is used on datasets from across different life stages, and on several populations from across a large portion of the natural geographical range of a single host species, revealing new details about the ecology and evolution of interactions between *M. cinxia* butterflies and their diverse parasites.

Previous studies have identified the butterfly *M. cinxia* as the host of several parasitoid species, including specialist and generalist parasitoids attacking the eggs, larvae or pupae of *M. cinxia* (Lei et al. 1997b; van Nouhuys 2016). We present evidence that some of our *M. cinxia* larvae were parasitized by either Braconidae or Ichneumonidae parasitoid wasps, most likely *Cotesia melitaearum* and *Hyposoter horticola*, which are both common in the Åland Islands (van Nouhuys 2016; van Nouhuys et al. 2012). The detected parasitism rates, however, (12% of our larval samples) remain under the expected 30% emergence rate by *H. horticola* alone in the Åland population (Duplouy et al. 2015; Montovan et al. 2015). Although, it is possible that this result denotes the low detection ability of the screening method, this result might still suggest that the parasitoids indeed only parasitize a 30% fraction of the available larval hosts (Montovan et al. 2015), rather than a much higher rate but experiencing some degree of mortality before emergence. Additionally, the lack of detection of parasitoid DNA from the sequenced adult butterflies is concordant with the idea that *Hyposoter* and *Cotesia* parasitoids are specialists of the *M. cinxia* larval stage, and that parasitism is fatal for the larvae (Montovan et al. 2015).

Our results provide the first evidence that *M. cinxia* is host to bacterial endosymbionts, including *Spiroplasma* and *Wolbachia*. *Spiroplasma* is a common symbiont of insects, infecting about 10% of species worldwide (Duplouy & Hornett 2018). The bacterium induces diverse phenotypes in its hosts (Jiggins et al. 2000; Pollmann et al. 2022), but its effects on the ecology and evolution of *M. cinxia* remain unknown (Duplouy et al. 2020a). As *Spiroplasma* was found at an intermediate prevalence (i.e. 45/169 samples, 26.6%) and conserved in all populations screened, we suggest that the symbiont may be of relevance for the fitness and/or ecology of its *M. cinxia* host. Several previous microbiota studies focusing on the Åland population have failed to highlight the presence of the symbiont (Duplouy et al. 2018; Duplouy et al. 2020a; Minard et al. 2022; Minard et al. 2019). We suggest that *Spiroplasma* is present at such low densities in larval and adult *M. cinxia* hosts, that its detection through metabarcoding approaches might not always be successful. This hypothesis is supported by the fact that, even after pooling all our samples, we could only produce a short partial *Spiroplasma* assembly (65KB) with very low read coverage. Future studies specifically investigating *Spiroplasma* in *M. cinxia* are needed to clarify the prevalence, titer, tropism and role of the symbiont in this butterfly species. In contrast, *Wolbachia*, perhaps the most common symbiont in insects worldwide (Duplouy & Hornett 2018; O’Neill et al. 1992; Zug & Hammerstein 2012), was rarely detected in *M. cinxia*. *Spiroplasma* and *Wolbachia* can sometimes co-infect their hosts (Mathé-Hubert et al. 2019), however, as both symbionts are thought to compete for the same resources, they might also exclude each other, which would explain the high prevalence of *Spiroplasma* and the low prevalence of *Wolbachia* in our system.

*Wolbachia* and *Spiroplasma* symbionts are predominantly transmitted maternally, but evidence of horizontal transfers of the symbionts is available from other host systems (Brown & Lloyd 2015; Duplouy et al. 2020b; Vavre et al. 1999). Parasitoid species in particular are described as potential vectors of *Wolbachia* to naïve hosts (Vavre et al. 1999). In the Åland Islands, 50% of the population of *H. horticola* is infected with a strain of the bacterial symbiont *Wolbachia* (Duplouy et al. 2015; Duplouy et al. 2021; van Nouhuys et al. 2016). The lack of infection in these particular wasps (Duplouy et al. 2015; Duplouy et al. 2021), or alternatively, the low sequencing coverage of our samples, combined with the possibly low density of *Wolbachia* in young parasitoid larvae, could explain why this *Wolbachia* strain was not detected in any of the six Finnish larvae parasitized by *Hyposoter*. Unfortunately, because of low coverage, we could not compare the partial *Wolbachia* assembly from the Spanish and Swiss *M. cinxia* samples to previously sequenced material from the *Wolbachia* strains infecting *H. horticola* from Åland or France (Duplouy et al. 2015). In contrast, we have no evidence of *Spiroplasma* infection in parasitoids of *M. cinxia* (Valerio et al. 2024b).

Overall, the analysis of metagenomes from different life-stages and populations revealed details of the interactions between the butterfly *M. cinxia* and its parasitoids, as well as with various symbiotic species. Although this study demonstrates the power of such analyses, applicable to any study system (Vancaester & Blaxter 2023), they also generate several new questions and avenues for further research on species interactions. For example, in addition to *Wolbachia* and *Spiroplasma*, other microbial infections, such as *Yersinia* bacteria and baculoviruses, were detected. The detection of such possibly pathogenic species is exciting, and their possible role in host population decline (Kahilainen et al. 2018) deserves further investigation.

Finally, although the development of more affordable whole-genome sequencing methods now supports the wider use of those methods in evolutionary biology, the overall coverage and total number of reads obtained from these sequencing projects may often be too low to allow the comprehensive and reliable detection of infections in the targeted host. In our case, even after merging individual sequencing files, we were only able to produce partial assemblies of the genomes of some of the symbiotic strains present in the different populations of *M. cinxia*. Furthermore, because of the overall low coverage of our samples, we remain cautious about some of our results. For example, the estimated prevalence of many infections should be interpreted carefully, as especially false negatives might still occur despite merging of the data. Similarly, because of the merging of specimen data, we cannot tell exactly how many individual specimens were truly infected per population. Nonetheless, as the outputs from two independent tools, Blobtool (Laetsch & Blaxter 2017) and CC-Metagen, (Marcelino et al. 2020), converged in their taxonomic assignments of the non-host genomic material, we are confident in the provided genus-level taxonomic identifications. But higher coverage would have provided us with better quality assemblies of all species, and would have provided additional population genetic information on the different parasites (Kankare et al. 2005).

In conclusion, even if represented at low coverage, available whole-genome sequencing projects not only provide the research community with genomic material from the target species, but also allow the discovery and potential monitoring of diverse pathogenic, parasitic, and symbiotic organisms associated with the target host. In the future, the results of our metagenomic analyses will guide the design of target experimental assays to reveal the role of each detected infection on the fitness and population dynamics of *M. cinxia*, as well as their importance for the butterfly’s ecology and evolutionary trajectory.

## Data availability

Supplementary material (Table S1-S3 and genomic assemblies) can be found in Zenodo under the DOI <u>10.5281/zenodo.14673566</u>. Raw sequence reads from the three re-sequencing datasets will only be made publicly available in NCBI upon publication of the research.

## Acknowledgments

We thank F. Valerio for providing reference genome databases for *Wolbachia*, *Spiroplasma* and *Cotesia*. The project was funded by the Academy of Finland (grant #321543 to AD, #316227 to MFD) and Helsinki Institute of Life Science (HiLIFE) start up package to M.S., M.M.J. and P.R., as members of the University of Helsinki Biodata Analytics Unit, were supported by the Academy of Finland’s ‘Thriving Nature’ research profiling action. Thank you to Constanti Stefanescu, Alma Oksanen, Debra DiLeo, Sandra Fagan, Antoni Mariné, and Rachel de Marco for assistance with sample collection in Spain, Switzerland, and Morocco.

## Supplementary material

**Figure S1:**
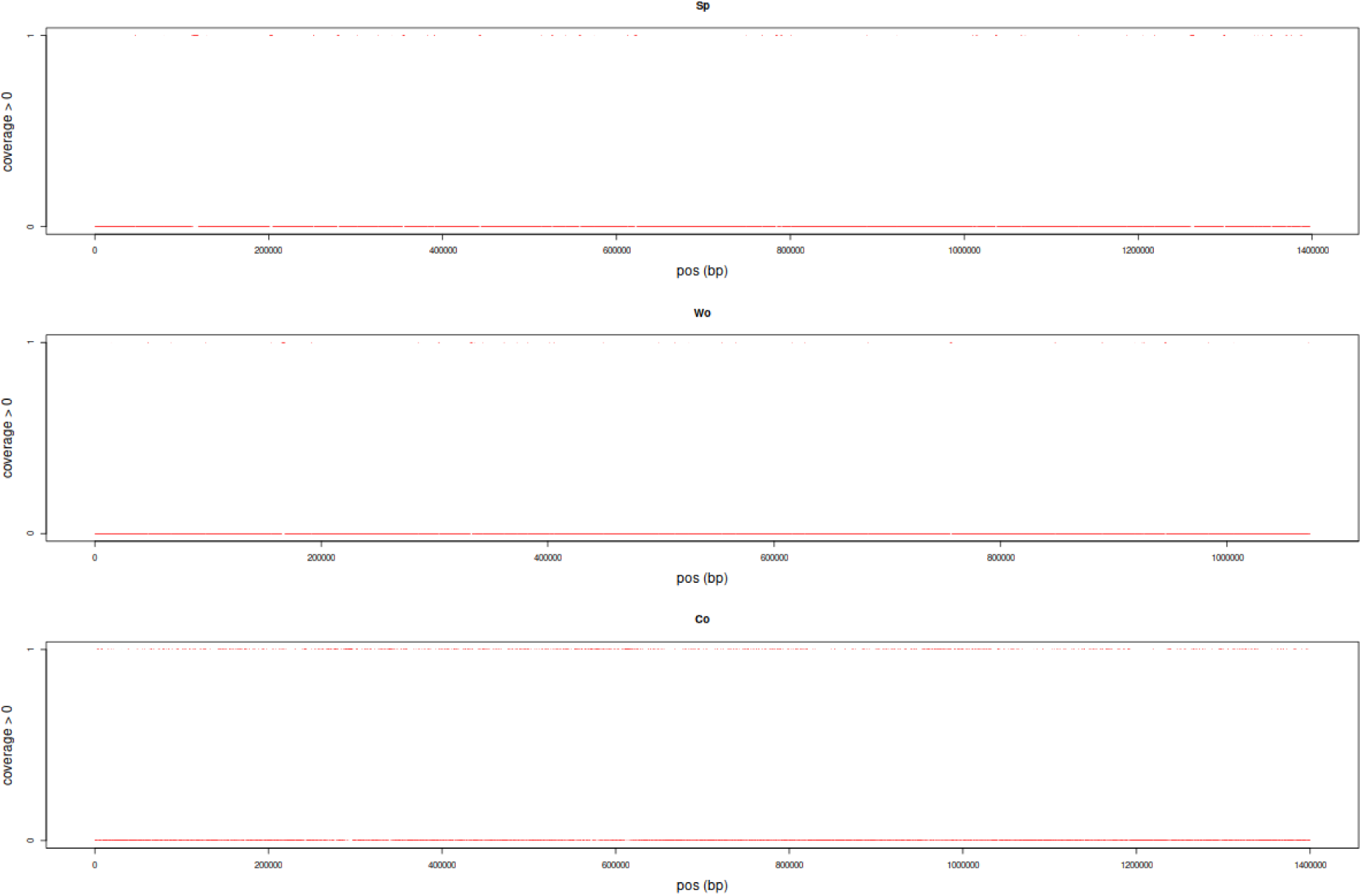
The coverage of *Spiroplasma* genome (top), *Wolbachia* genome (middle), and the first 1.4Mb of first *Cotesia* chromosome (bottom). The top row (coverage=1) of each graph shows the low coverage across the chromosomes. Only very few positions have been covered explaining the resulting partial assemblies obtained in the present study.

**Table S1: Outputs from the Blobtool analyses.** Data can be found under Zenodo DOI <u>10.5281/zenodo.14673566</u>.

**Table S2: Number of reads assigned to each of three parasites (ie. *Cotesia, Spiroplasma, Wolbachia*) for each sample.** Data can be found under Zenodo DOI <u>10.5281/zenodo.14673566</u>.

**Table S3: Output from the CCMetagen analyses on five selected samples.** Data can be found under Zenodo DOI <u>10.5281/zenodo.14673566</u>.

